# B-SIDER: Computational Algorithm for the Design of Complementary β-sheet Sequences

**DOI:** 10.1101/631069

**Authors:** Tae-Geun Yu, Hak-Sung Kim, Yoonjoo Choi

## Abstract

The β-sheet is an element of protein secondary structure, and intra-/inter-molecular β-sheet interactions play pivotal roles in biological regulatory processes including scaffolding, transporting, and oligomerization. In nature, a β-sheet formation is tightly regulated because dysregulated β-stacking often leads to severe diseases such as Alzheimer’s, Parkinson’s, systemic amyloidosis, or diabetes. Thus, the identification of intrinsic β-sheet forming propensities can provide valuable insight into protein designs for the development of novel therapeutics. However, structure-based design methods may not be generally applicable to such amyloidogenic peptides mainly owing to high structural plasticity and complexity. Therefore, an alternative design strategy based on complementary sequence information is of significant importance. Herein, we developed a database search method called B-SIDER for the design of complementary β-strands. This method makes use of the structural database information and generates query-specific score matrices. The discriminatory power of the B-SIDER score function was tested on representative amyloidogenic peptide substructures against a sequence-based score matrix (PASTA2.0) and two popular *ab initio* protein design score functions (Rosetta and FoldX). B-SIDER is able to distinguish wild-type amyloidogenic β-strands as favored interactions in a more consistent manner than other methods. B-SIDER was prospectively applied to the design of complementary β-strands for a splitGFP scaffold. Three variants were identified to have stronger interactions than the original sequence selected through a directed evolution, emitting higher fluorescence intensities. Our results indicate that B-SIDER can be applicable to the design of other β-strands, assisting in the development of therapeutics against disease-related amyloidogenic peptides.

## Introduction

The β-sheet is one of the major units of protein structure ^1^, and has a variety of functions in transportation, recognition, scaffolding, and enzymatic processes ^2^. The mechanism of a β-sheet formation has recently received significant attention owing to its close relations with several critical diseases such as Alzheimer’s, Parkinson’s, type 2 diabetes, and systemic amyloidosis ^3, 4^. Such diseases are known to be linked to the precipitation of dysregulated β-stacking between neighboring β-strands ^5-7^. In this regard, information on the amino acid propensity of intrinsic β-sheet forming motifs and its use in the design of their complementary sequences are crucial for understanding the mechanism of β-sheet formation and developing potential therapeutics specifically targeting aggregation-prone regions ^8^.

Whereas structure-based protein design approaches have shown notable successes in several cases ^9^, their application to *de novo* β-sheet designs still remains a challenge ^2, 10^. Although structure-based design approaches require a well-defined protein structure, amyloidogenic peptides usually have highly disordered structures ^11^. The structural identification of such peptides has long been hindered by high degrees of structural plasticity, transiency, and complexity owing to a self-oligomerization ^12, 13^. It is thus necessary to exploit the complementarity across neighboring β-strand pairs using sequence information.

Intriguingly, significant conservation and covariations of residue pairs between neighboring β-strands have been identified in many different protein families ^14^. For instance, pairs of β-branched residues and cysteines are preferred at nonhydrogen-bonded positions. Aromatic residues tend to be paired with valine or glycine ^15^. Several computational algorithms have been developed to predict aggregation-prone regions based on the internal β-sheet forming patterns. While differing in detail, they make use of either statistical potentials such as Tango ^16^, PASTA ^17^, SALSA ^18^, BETASCAN ^19^, and Waltz ^20^ or physicochemical properties of amino acids ^21^. In addition, consensus methods and machine-learning approaches have also been developed ^22, 23^, showing fine agreement with the experiment results.

It has been reported that β-strand interactions can be stabilized by introducing β-sheet favored pairs ^24-28^ and charge pairing between neighboring β-strands ^29-31^. Recent studies have shown that fragments derived from the amyloidogenic region can be used for β-stack modeling ^6, 32^. While the use of amino acid pairing information in a protein design has been attempted elsewhere, practical applications of such patterns have been limited, mainly owing to the lack of a comprehensive quantification for residue pairing and noisy patterns of β-sheet forming residue pairs ^1, 33, 34^. The β-sheet forming peptides appear to have poor sequence commonalities and imperfect repeats ^19^. Therefore, a careful curation of meaningful patterns is required for a practical protein design strategy of complementary β-strands.

Herein, we present a database search method, B-SIDER (β-Stacking Interaction DEsign for Reciprocity), for the design of complementary β-strands. The method generates a query-specific score matrix from the structure database. To utilize the pairing information and overcome the pattern noise, we hypothesized that significant complementary pairs can be amplified by superposing a subset of sequence fragments. Moreover, the recent growth boom of β-sheet structures ^35^ allows the solid statistics of β-sheet forming residue pairings ^36^. Based on the hypothesis and statistics of β-sheet forming residue pairings, we developed a fast and reliable computational method for the design of complementary β-strand sequences. The methodology augments β-sheet forming residue preferences by overlaying complementary fragment sequences (**Figure 1**). We retrospectively validated our approach using a set of curated amyloidogenic targets and compared it with two popularly used structure-based methods (Rosetta and FoldX) and a sequence-based aggregation prediction algorithm (PASTA 2.0). Our algorithm was shown to clearly distinguish favorable β-sheet forming sequences entirely based on the query sequence, whereas the structure-based energy functions exhibited inconsistent results depending on the targets. The utility and potential of our method were demonstrated by designing novel complementary peptides for splitGFP. The designed sequences showed stronger interactions with neighboring strands of the scaffold, and consequently higher fluorescence emissions, than the original peptide selected through directed mutagenesis ^37^.

**Figure 1.**
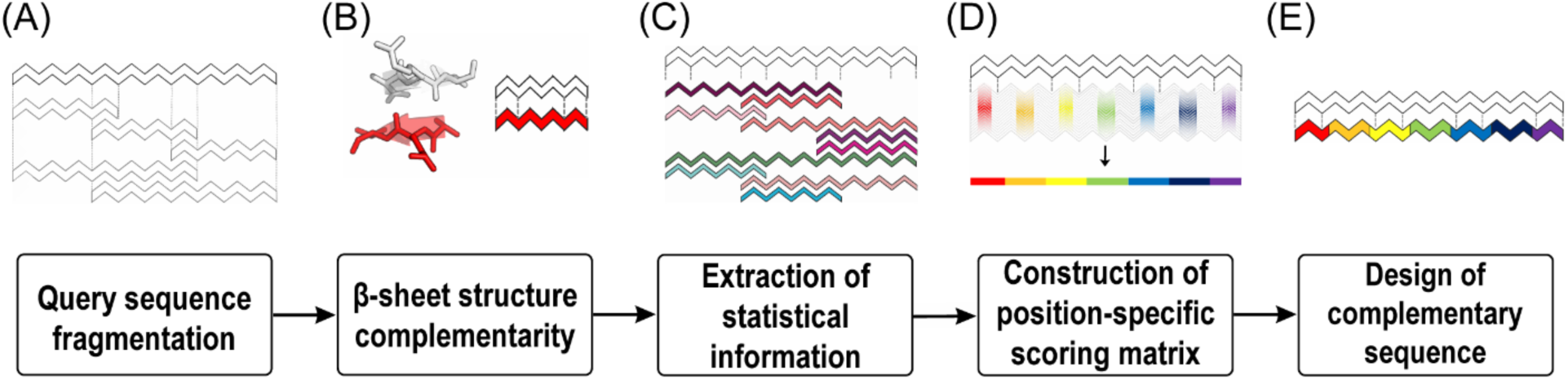
Overview of the B-SIDER algorithm. (A) When a query sequence is given, the query is divided into smaller linear peptides ranging from three residues to its full length. (B) Exactly matched sequences are identified and checked against the structure database such that the matches indeed form β-sheets. (C) If the matches form β-sheets, their complementary sequences are extracted. (D) A position-specific score matrix is constructed based on the complementary sequence information. (E) Final complementary sequences are designed using the score matrix.

## Materials and Methods

### Computational algorithm for the design of complementary sequences

#### Collection of β-strand information

Non-redundant structures determined by high-resolution X-ray crystallography were collected from the PISCES ^38^ (10∼90% sequence identity), < 3 Å resolution, < 0.3 R-factor and sequence length from 40 to 10000. Given the query target sequence, the non-redundant structure database was used to extract pairing information from matched sequences with the same directionality. The target sequence is initially divided into linear moving-windows whose residues in length range from 3 to the entire target-sequence length. Any structures with identical target sequence fragments as the split queries were collected, followed by further filtering based on the definition of β-sheet secondary structure (the distance between the backbone nitrogen-oxygen atom pairs < 5 Å). To remove any redundancy, protein structures that contain matches were compared using TMalign ^39^. If the TM-score > 0.7 and sequence identity > 90%, one of the matched sequences is removed.

While the method is applicable to both parallel and anti-parallel β-sheets in theory, we entirely focused on anti-parallel β-sheets in this study ^27, 28^ and all the test cases are anti-parallel because anti-parallel cases are more frequently observed compared to parallel cases ^40^. Disease-related amyloidogenesis is also known to be initiated with anti-parallel β-sheets, and soluble oligomeric amyloid species mainly exist as anti-parallel ^41, 42^.

#### Complementary sequence score

The β-sheet complementarity score function is derived from the environment-specific substitution score ^43^. We hypothesized that each position of a β-strand is independent of one another, and their complementarities are determined through residue pairs from neighboring strands. Given the query sequence, all identified neighboring sequences are pooled together, as described in the previous section. The amino acid frequency at each complementary position is counted as

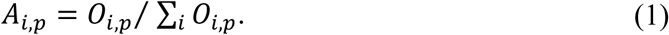

where *A*_*i,p*_ is the frequency of a certain amino acid (*i*) at a specific complementary position (*p*), and *O*_*i,p*_ is the total count of the amino acid at *p*. The background frequency of a certain amino acid *B*_*i*_ is independent of the query and thus counted from a well defined structure database(HOMSTRAD database ^44^). The frequency is calculated as

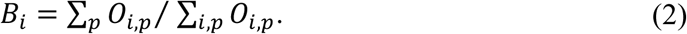

The complementary sequence score of the amino acid at the position *S*_*i,p*_ is calculated as

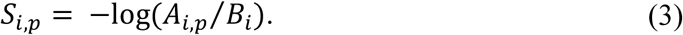

It should be noted that the complementarity score is completely data-driven, i.e., if an amino acid never appears at a certain position, a high penalty score is imposed. We only consider complementary amino acids that are found at least once in the entire identified sequences. The final score is the sum of the scores at all positions.

### Protein expression and complementation assay

#### Gene construction

The gene coding for splitGFP ^37^ consists of the first through tenth (GFP1-10) and 11^th^ strand (GFP11) templates (Supporting Information, **Table S1**). They were cloned into pET-28a (Novagen) vector between the Nde-I and Xho-I restriction sites. We introduced additional mutations to GFP1-10 to inhibit aggregation and for a convenient expression ^45^. The GFP11 strand was fused with a P22 virus-like particle scaffolding protein ^46^ for a soluble and stable expression. Mutations on GFP11 were introduced through PCR using mutagenic primers (Supporting Information, **Table S2**), and the resulting genes were cloned into the pET-28a vector. Six histidine residues were fused to the N-terminal of the GFP1-10 and GFP11 genes as an affinity purification tag.

#### Protein expression and purification

All constructs were transformed into BL21 (DE3) Escherichia coli strains. The transformed cells were grown overnight and inoculated into a Luria-Bertani media containing 50 µg/ml of kanamycin at 37 °C. The cells were then grown until the optical density of the cells reached 0.6–0.8 at 600 nm, followed by the addition of 0.7 mM of IPTG (isopropyl β-D-1-thiogalactopyranoside) for induction. After incubation for 16–18 h at 18 °C, the cells were harvested and suspended in a lysis buffer containing 50 mM Tris (pH 8.0), 150 mM NaCl, and 5 mM imidazole. The suspended cells were disrupted through sonication, and insoluble fractions were removed using centrifugation at 18,000 g for 1 h. The supernatants were filtered with 0.22 µm syringe filters and purified through affinity chromatography using Ni-NTA agarose Superflow (Qiagen). The solutions were applied to the resin-packed columns and washed with a buffer containing 50 mM Tris (pH 8.0), 150 mM NaCl, and 10 mM imidazole, until no proteins were detected through a Bradford assay. An elution buffer (50 mM Tris (pH 7.4), 150 mM NaCl, 300 mM imidazole) was then applied to the column. The buffer exchange was performed with phosphate buffered saline (PBS, pH 7.4) using a PD-10 column (GE health-care). The concentrations of the proteins were determined by measuring the absorbance at 280 nm. All purification processes were conducted at 4 °C. The purities of the proteins were then evaluated using SDS-PAGE.

#### Complementation assay

The assembly of splitGFP variants was monitored and measured using a fluorescence complementation assay. An excessive amount of GFP1-10 (50 pmol) in 180 µl and 20 µl of equal molar concentration for each GFP 11 strand (3 pmol) were co-incubated in a PBS buffer (pH 7.4). The fluorescence kinetics (λ_ex_= 488 nm /λ_em_= 530 nm) were monitored for 12 h at 25 °C using a TECAN infinite M200 microplate reader at 5 min intervals ^37^ with shaking for 2 sec between intervals. Each experiment was conducted in triplicate using a Nunc F 96 Micro-well black plate, blocked with a solution of PBS containing 0.5 % of bovine serum albumin (BSA) for 30 min before the assay.

## Results/Discussion

### Overview of the design process

We hypothesized that repetitively observed amino acid pairing patterns indicate a strong preference toward the β-sheet. It was also assumed that the sequence with the most frequent patterns will directly form a β-sheet without considering other environmental contributions.

There are two major steps applied in the algorithm: 1) The extraction of complementarity information of the β-sheet and 2) the construction of a scoring matrix (**Figure 1**). When a query sequence is given, it is fragmented into several pieces of short peptides longer than three residues in length, and the matching neighboring strands are collected. These fragmentation and overlaying processes naturally impose weights on complementary-prone positions and amplify the pattern signals. It should be noted that the subsequence search starts from the −2^nd^ position in order to avoid underweighting teminal regions (Supporting Information, **Figure S1**). After the collection of matched sequences, a position-specific complementarity scoring matrix is constructed. The scoring matrix obtained is used to evaluate and design the complementarity of β-strand interactions.

### Validation of the score function on retrospective cases

In an effort to validate the complementarity score, we manually curated a test set of naturally occurring β-strand pairs whose environmental effects are minimal. It is known that β-strand pairing is in general significantly hindered by local environments ^47^, whereas amyloidogenic peptide segments are known to form natural β-sheets themselves ^17^. We thus selected a set of widely known amyloidogenic structures whose aggregation-prone regions have been identified (**Table1** and Supporting Information, **Table S3**). When multiple β-strands are present, we assumed that the first strands may be the primary amyloid building unit and so only the first two strands as a pair were extracted for the structure-based energy calculation.

**Table 1.**
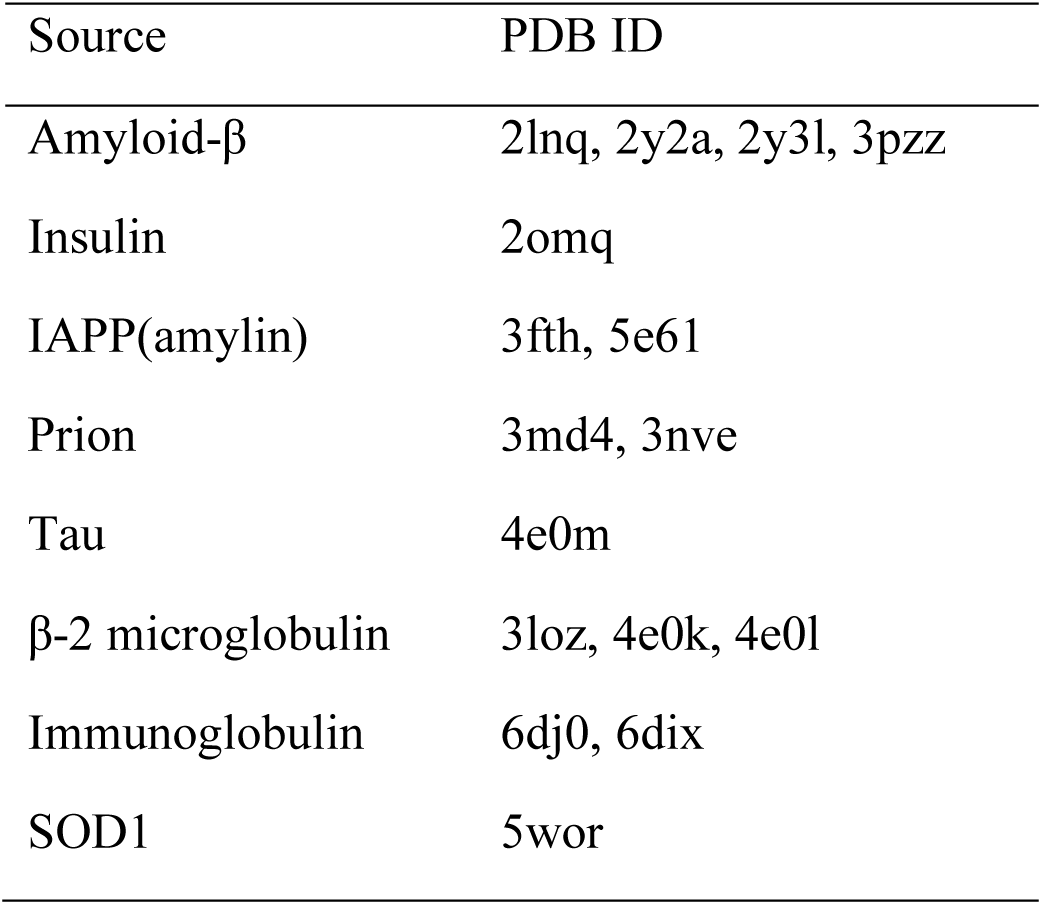
Retrospective test set.

To assess the complementarity of the native sequences, we compared their scores with those of random sequences which were generated using all 20 amino acids in a position-independent manner. Natural amyloidogenic segments are known to be highly prone to aggregation, and thus they are expected to be highly preferred, i.e., having fairly low scores in the random sequence score distributions. **Figure 2** shows that all native sequence scores are ranked extremely low in all distributions. On average, the native sequences are within 5.2% of the distributions (**Figure 2**). The results indicate that the scoring function is extremely useful in detecting favorable β-strand counterparts.

**Figure. 2.**
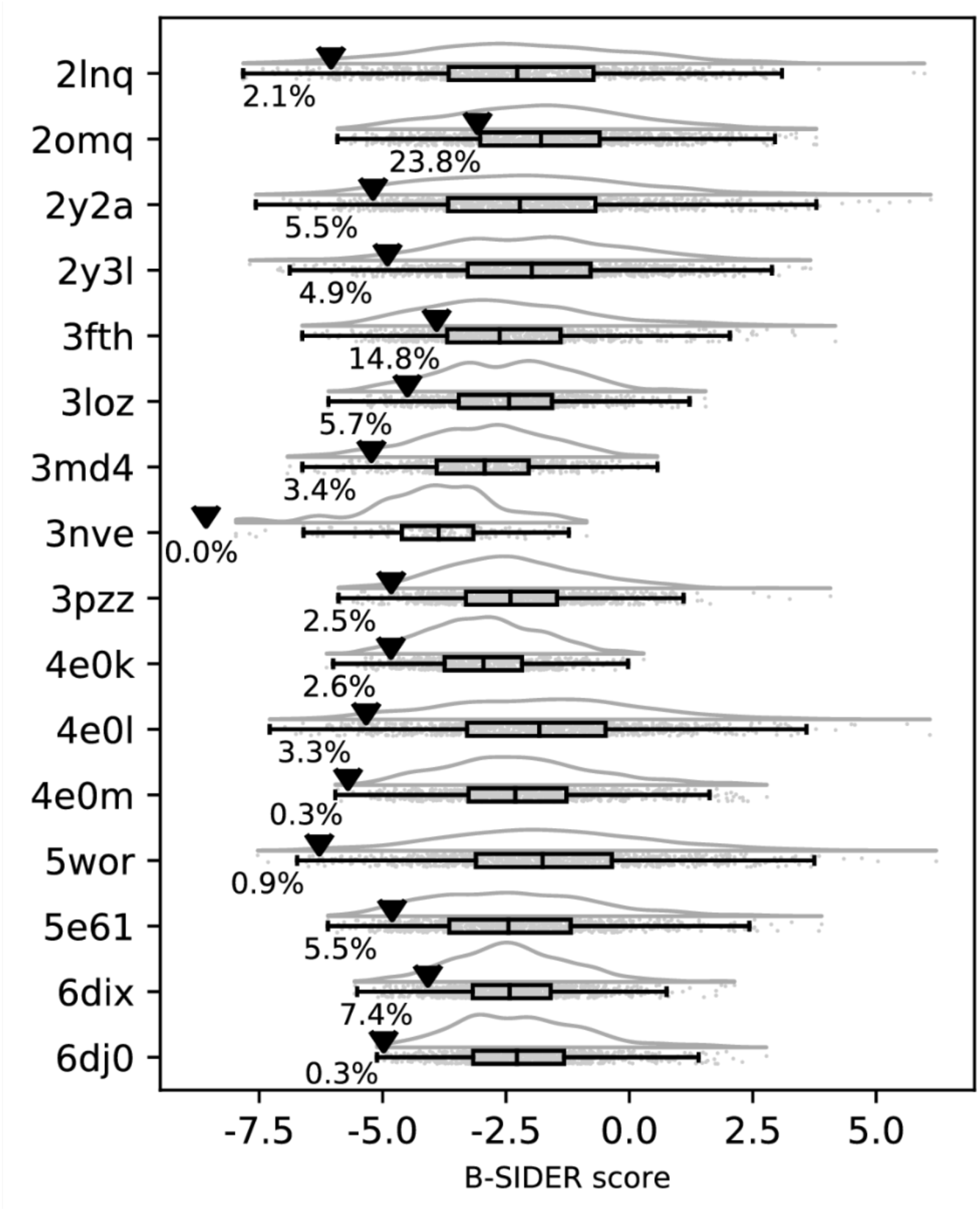
The predictive power of the B-SIDER score. In this test set, native sequences were compared against 1,000 random sequences. A lower score indicates more favorability. The native complementarity scores are marked as (▾) and their percentile values are displayed next to the marks. This plot was generated using the Raincloud Python package ^48^.

We also compared the B-SIDER score with two structure-based all-atom energy functions and a sequence-based score matrix. For the structure-based methods, we picked Rosetta (Talaris 13) ^49, 50^ and FoldX ^51^, which have been popularly used in *de novo* protein designs ^52, 53^. PASTA 2.0 ^54^ is a method for predicting regions prone to aggregation using a scoring matrix derived from residue pairing patterns of β-sheets. To avoid any biases, 1,000 random sequences were newly prepared per target. The “FastRelax” protocol ^55^ from Rosetta (Ver. 3.7), the “BuildModel” command from FoldX (Ver. 4.0), and the scoring matrix from PASTA 2.0 were used against the native and random sequences. The predictive power of the score function was assessed based on the percentile of the native sequence score against the random sequence score distribution.

**Figure 3A** shows that structure-based score functions are generally worse than the sequence-based scoring matrices. The results indicate that the Rosetta energy score function is not sufficiently accurate for ranking complementary β-strands (35.1^th^ percentile on average), whereas the predictive powers of PASTA 2.0 and FoldX were moderate, showing 7.4^th^ and 16.4^th^ percentiles on average respectively. B-SIDER was shown to be the most accurate in an extremely consistent manner (5.2^th^ on average. Standard deviations of Rosetta, FoldX, PASTA 2.0 and B-SIDER are 25.2, 23.3, 8.7 and 6.2, respectively). Although the assessment by PASTA 2.0 is also fairly consistent, the query-specific nature of B-SIDER may provide better results.

**Figure. 3.**
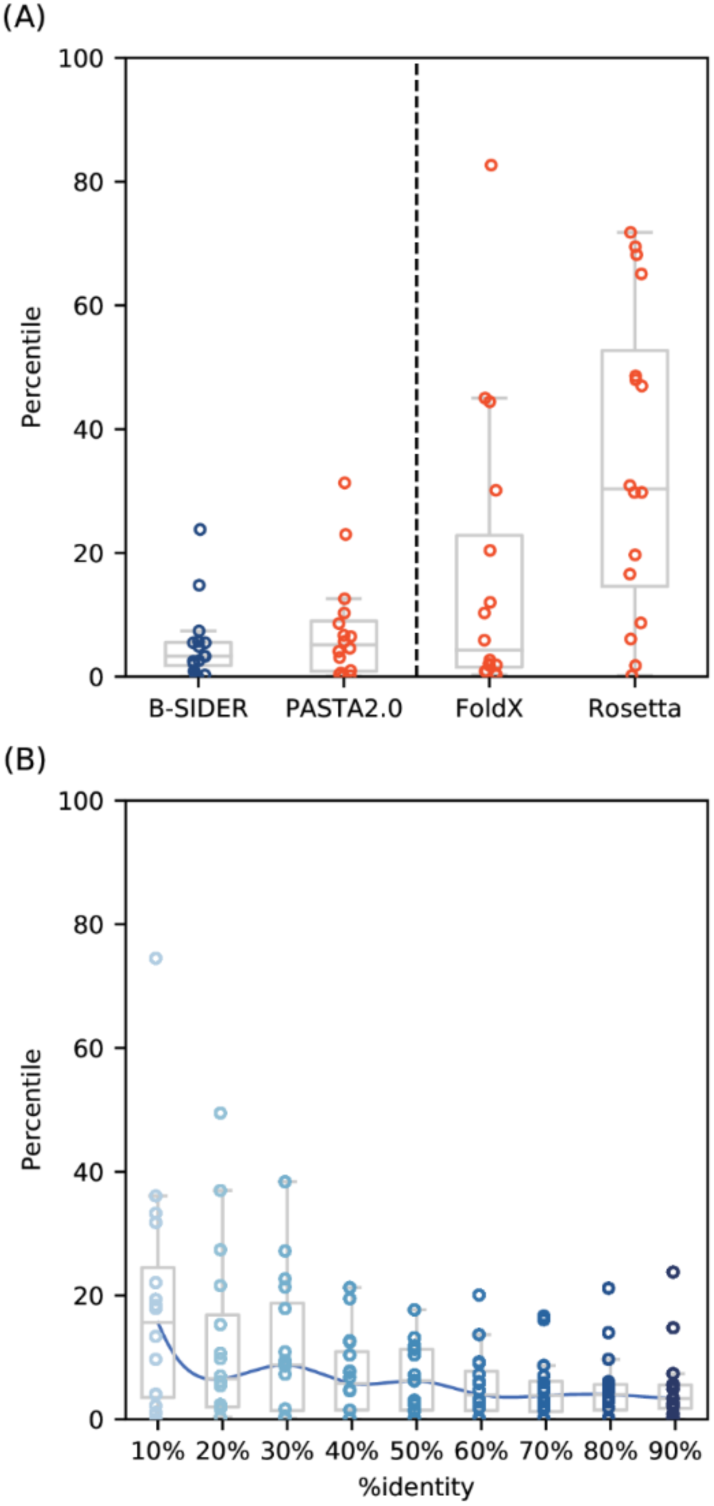
Predictive power of B-SIDER score and its robustness. (A) The B-SIDER score is compared against two popularly used structure-based protein design methods (Rosetta and FoldX) and the scoring matrix for the prediction of an amyloid formation (PASTA 2.0). For a fair comparison, we generated new sets of 1,000 random sequences per target and assessed their scores using each energy function. The percentile values of native complementary sequences against the score distributions of the randomly generated sequences are presented. The B-SIDER score provided the most consistent assessment. (B) The robustness of the B-SIDER score was evaluated at various sequence identity threshold values. The percentile values of B-SIDER are overall fairly consistent at even low sequence identity cutoff.

Considering that the Rosetta relax protocol performs flexible backbone refinements, the use of a fixed-backbone calculation seems to be better for an evaluation of the β-sheet complementarity. It should be noted that the inconsistent results of Rosetta imply that FoldX prediction will also be highly driven by the preparation of the structure, i.e., a design with an ill-defined model may not be generally successful. By contrast, B-SIDER and PASTA 2.0 do not require any structure at all, and thus can be applied to general cases such as β-sheet interactions with high structural plasticity and poor structural integrity, which are the common features of amyloidogenic peptides. Furthermore, the process of collecting complementary motifs of B-SIDER also appears to be powerful, making it possible to distinguish favorable complementary sequences not easily detected through one-to-one residue pairing. We also tested the robustness of the B-SIDER approach by examining the algorithm at various sequence identity cutoff values of the structure database. The results are fairly consistent until < 40% sequence identity cutoff (**Figure 3B** and Supporting Information, **Figure S2)**. We observe that significant outliers start appearing at < 30%. Despite such outliers, B-SIDER still successfully discriminates most test cases as favorable. The length distributions of matched subsequences (**Figure S3**) show that complementarity information is mainly derived from subsequences with the minimal length. Perhaps the consistency of the predictive power at low homology thresholds may come from the subsequence overlapping.

### Prospective appplication of the algorithm to splitGFP

As shown in the retrospective test, B-SIDER is extremely useful in discriminating naturally β-strand forming sequences. As a proof of concept, we prospectively designed novel complementary β-strands for splitGFP. SplitGFP is a fragmented protein pair derived from superfolderGFP ^37^, comprising a scaffold containing ten β-strands (GFP1-10) and their complementary β-strand peptide (GFP11). GFP11 specifically interacts with GFP1-10, and the strand tightly forms a stable β-sheet structure, which facilitates the chromophore maturation in an irreversible manner ^56^. This assembly process results in the emission of green fluorescence. Because GFP11 is known to be disordered in a solution, its conformational transition from a disordered to an induced β-sheet is similar to that of amyloidogenic peptides ^57, 58^. This model system thus efficiently assesses whether sequences designed using B-SIDER have favorable β-sheet interactions.

The original GFP11 was designed using directed mutagenesis, and it shows a high intrinsic propensity to form hydrogen bonds with the neighboring β-strands of GFP1-10 ^59^. In our case, the queries are the neighboring strands of GFP1-10 (**Figure 4A**). The total score was calculated as a sum of the scores from both sides without specific weights. It is known that the inward pointing residues (1, 3, 5, and 7^th^ positions) directly interact with the chromophore, and thus are not subject to a mutation. It should be noted that we utilized the best structure data available (PDB < 90 % sequence identity). B-SIDER identified 2,637 non-redundant sequences from the structure database. The native sequence is ranked modestly among 1,000 possible randomly chosen sequence variants (46^th^ percentile), indicating that there could be room for complementary sequences with stronger interactions than the original (**Figure 4B**). We then selected sequences with the lowest B-SIDER scores (top_vars; **Table2**). Amino acid compositions of the ten variants are mostly hydrophobic or branched amino acids (Supporting Information, **Figure S4**). In addition, one sequence with a high score (> 75^th^ percentile) was randomly selected as a negative control (neg_var, 77^th^).

**Figure. 4.**
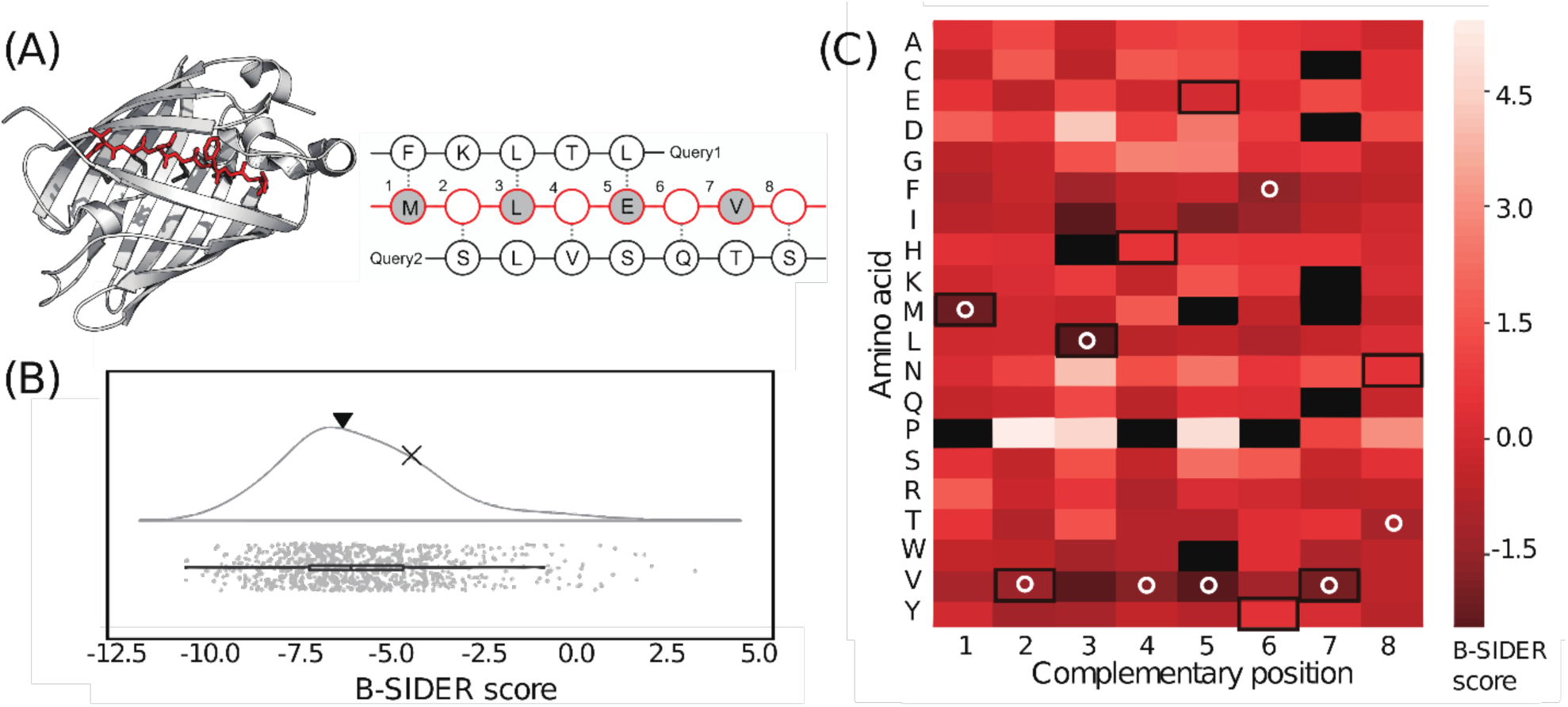
Design process of GFP11 variants. (A) The strand to be designed (GFP11) is highlighted in red, and query sequences to consider are shown on the right. Dotted lines represent hydrogen bonds. The residues pointing inward, which are not subject to a mutation, are colored in gray. (B) The B-SIDER score of the original GFP11 (▾) is compared against the score distribution of randomly generated sequences. The negative variant (neg_var) is marked as × (the 77^th^ percentile). (C) The position-specific scoring matrix for the query sequences. Amino acids that never appeared at each position are colored in black. The amino acids of the native sequence are highlighted with black boxes. The white circles represent the lowest scores at each position.

**Table 2.**
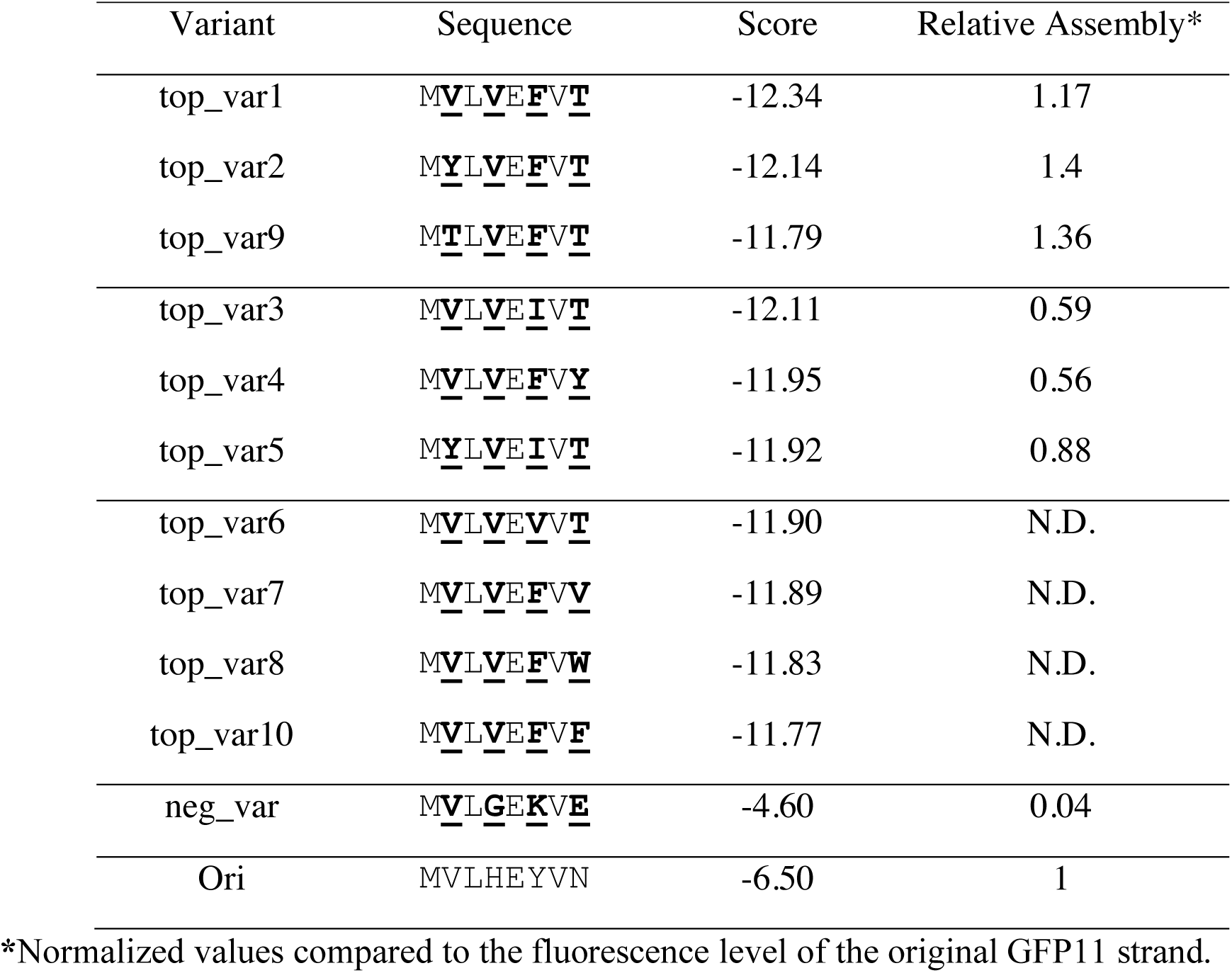
GFP 11 variants.

| Variant | Sequence | Score | Relative Assembly* |
| --- | --- | --- | --- |
| top_var1 | M<u>V</u>L<u>V</u>E<u>F</u>V<u>T</u> | -12.34 | 1.17 |
| top_var2 | M<u>Y</u>L<u>V</u>E<u>F</u>V<u>T</u> | -12.14 | 1.4 |
| top_var9 | M<u>T</u>L<u>V</u>E<u>F</u>V<u>T</u> | -11.79 | 1.36 |
| top_var3 | M<u>V</u>L<u>V</u>E<u>I</u>V<u>T</u> | -12.11 | 0.59 |
| top_var4 | M<u>V</u>L<u>V</u>E<u>F</u>V<u>Y</u> | -11.95 | 0.56 |
| top_var5 | M<u>Y</u>L<u>V</u>E<u>I</u>V<u>T</u> | -11.92 | 0.88 |
| top_var6 | M<u>V</u>L<u>V</u>E<u>V</u>V<u>T</u> | -11.90 | N.D. |
| top_var7 | M<u>V</u>L<u>V</u>E<u>F</u>V<u>V</u> | -11.89 | N.D. |
| top_var8 | M<u>V</u>L<u>V</u>E<u>F</u>V<u>W</u> | -11.83 | N.D. |
| top_var10 | M<u>V</u>L<u>V</u>E<u>F</u>V<u>F</u> | -11.77 | N.D. |
| neg_var | M<u>V</u>L<u>G</u>E<u>K</u>V<u>E</u> | -4.60 | 0.04 |
| Ori | MVLHEYVN | -6.50 | 1 |

\*Normalized values compared to the fluorescence level of the original GFP11 strand.

The selected variants were successfully expressed and purified (Supporting Information, **Figure S5**) except for four clones (top_var6, 7, 8, and 10), which were observed to be insoluble, perhaps due to aggregation. Among those expressed, three variants (top_var1, 2, and 9) showed faster assembly patterns and higher signals compared to the original GFP11 (**Figure 5**). No functional aberrance with excitations or emissions was observed (Supporting Information, **Figure S6**). All successful variants, which emitted stronger fluorescence levels than the original, were shown to have pairs of phenylalanine and threonine at positions 6 and 8, respectively. These results demonstrate that the designed variants indeed formed complementary β-strands in a more favorable fashion than the original peptide as predicted. The other variants showed slightly lower signals than the original, but still gave rise to clear fluorescence signals (**Figure 5**). The negative control (neg_var) barely emitted any signal, suggesting that the score indeed indicates the complementarity of the β-stacking interactions. We also assigned scores to the GFP11 variants using other scoring methods. As shown in the retrospective test set, Rosetta and FoldX were unable to discriminate top_vars as favorable (Supporting Information, **Figure S7**). However, PASTA 2.0 was again fairly accurate in this case.

**Figure. 5.**
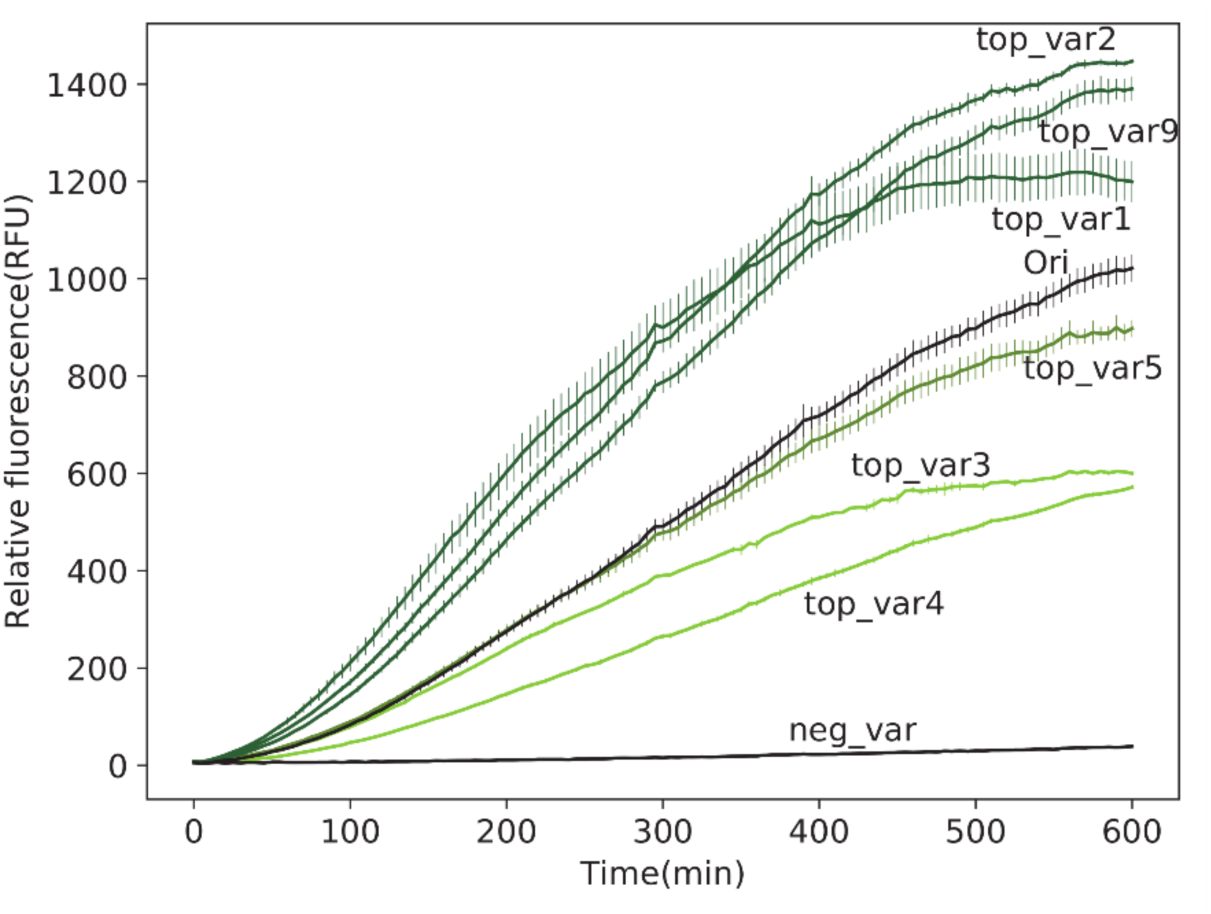
Complementation assay with the designed GFP11 variants. Among the six successfully expressed variants, three variants exhibited stronger fluorescence emissions than the original peptide identified through a directed mutagenesis.

## Conclusion

For understanding the aggregation mechanism of disease-related β-sheets and developing potential therapeutics against them, β-sheet forming patterns are essential. Unlike α-helices, however, there has been no established design principle for the complementarity of β-sheets. In this study, we developed B-SIDER, a database search method for the design of complementary β-strands based on the intrinsic β-sheet forming propensities. Statistical patterns of interacting residue pairs between neighboring β-strands enable the complementary interaction to be quantified. We demonstrated that the statistical potential can be directly applied to the design of complementary β-strand sequences. Using splitGFP as a model system, we successfully designed fragment variants, which led to stronger fluorescence emissions than the native one originally identified through a directed mutagenesis. The results clearly indicate that B-SIDER can be useful for the detection and design of β-stacking interactions between unstructured fragments. Therefore, our approach can find wide applications in protein designs where structure-based methods are ineffective, including the development of protein binders specifically against intrinsically disordered disease-related proteins.

## Notes

B-SIDER is freely available at http://bel.kaist.ac.kr/research/B-SIDER

## Supporting Information

Supporting Information Available: Sequences of GFP constructs (Table S1 and S2), identification of query sequences of the retrospective test set (Table S3), schematic description of B-SIDER scoring (Figure S1), discriminatory power of B-SIDER scoring at various sequence identity cutoff values (Figure S2), length distribution of matched subsequences (Figure S3), sequence logo of GFP11 variants (Figure S4), SDS-PAGE analysis of GFP11 variants (Figure S5), excitation/emission spectral analyses of GFP11 variants (Figure S6), score reassessment of the GFP11 variants using PASTA2.0, FoldX and Rosetta (Figure S7).

This material is available free of charge via the Internet at http://pubs.acs.org

## Acknowledgements

This work was conducted using Alphacom high-performance computing cluster at the department of biological sciences at the Korea Advanced Institute of Science and Technology (KAIST).

This work was supported by the Korea Research Fellowship Program (2016H1D3A1938246 to Y.C.), Global Research Laboratory (NRF-2015K1A1A2033346), and Mid-Career Researcher Program (NRF-2017R1A2A1A05001091) of the National Research Foundation (NRF) funded by the Ministry of Science and ICT.

## Abbreviations

GFP: green fluorescent protein;
PCR: polymerase chain reaction;
IPTG: isopropyl β-D-1-thiogalactopyranoside;
BSA: bovine serum albumin;
IAPP: islet amyloid polypeptide precursor;
SOD1: superoxide dismutase 1;
PDB: protein data bank

## For Table of Contents Use Only

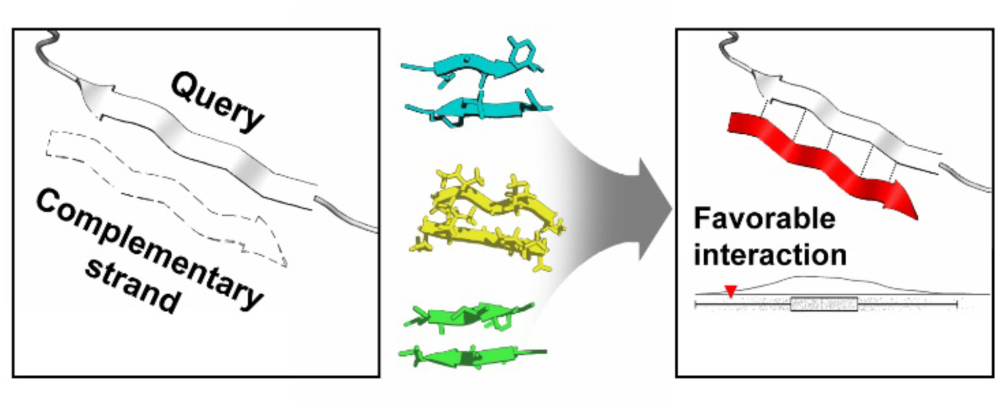

