## Supporting Information for "B-SIDER: Computational Algorithm for the Design of Complementary β-sheet Sequences"

**Table S1. Sequences of GFP1-10 and GFP11.** GFP11 (**bold**) and the designed region (**bold and underlined**) were fused with the P22 virus-like particle scaffold protein (*italic*)

| Name | Protein sequence | DNA sequence |
|---|---|---|
| GFP11 original | MGSSHHHHHHSSGLVPRGS**HMRDHM**<br>**VLHEYVN*AAGIT***GGSGGS*LTRLSER*<br>*LTLKPRGKQISSAPPADQPITGDVS*<br>*AANKDAIRKQMDAAASKGDVETYRK*<br>*LKAKLKGIR* | atgggcagcagccatcatcatcatcatcacagcagcggcctggtgccgcgcggcagc**c**<br>**atatgagagaccacatggtccttcacgagtacgtaaacgctgctgggattaca**ggc<br>ggttccggtggatct*ctcactcgactatccgaacgcttaactctcaagcctcgcggtaaa*<br>*caaatctcttccgctccccctgctgaccagcctattaccggtgatgtcagcgcagcaaa*<br>*taaagatgccattcgtaaacaaatggatgctgctgcgagcaagggagatgtggaaac*<br>*ctaccgcaagctaaaggcaaaacttaaaggaatccgataactcgag* |
| GFP1-10 | MGSSHHHHHHSSGLVPRGSHMKGE<br>ELFTGVVPILVELDGDVNGHEFSV<br>RGEGEGDATIGKLTLKFICTTGKL<br>PVPWPTLVTTLTYGVQCFSRYPDH<br>MKRHDFFKSAMPEGYVQERTISFK<br>DDGKYKTRAVVKFEGDTLVNRIEL<br>KGTDFKEDGNILGHKLEYNFNSHD<br>VYITADKQENGIKAEFTVRHNVED<br>GSVQLADHYQQNTPIGDGPVLLPD<br>NHYLSTQTVLSKDPNEK | atgggcagcagccatcatcatcatcatcacagcagcggcctggtgccgcgcggcagccat<br>atgaaaggagaagaacttttcactggagttgtcccaattcttgttgaattagatggtgatgttaa<br>tgggcacgaattttctgtccgtggagagggtgaaggtgatgcaacaatcggaaaacttaccc<br>ttaaatttatttgcactactggaaaactacctgttccatggccaacacttgtcactactctgactt<br>atggtgttcaatgcttttcccgttatccggatcacatgaaacggcatgactttttcaagagtgcc<br>atgcccgaaggttatgtacaggaacgcactatatctttcaaagatgacgggaaatacaagac<br>gcgtgctgtagtcaagtttgaaggtgatacccttgttaatcgtatcgagttaaaaggtactgatt<br>ttaaagaagatggaaacattctcggacacaaactcgaatacaactttaactcacacgatgtat<br>acatcacggcagacaaacaagaaaatggaatcaaagctgaattcactgttcgccacaacgtt<br>gaagatggctccgttcaactagcagaccattatcaacaaaatactccaattggcgatggccct<br>gtccttttaccagacaaccattacctgtcgacacaaactgttctttcgaaagatcccaacgaaa<br>agtaactcgag |

**Table S2. DNA Sequences of GFP11 variants.**

| Primer | Sequence |
|---|---|
| top_var1 | aataatcatatgagagaccacatggtgctggtggaatttgtgaccgctgctgg |
| top_var2 | aataatcatatgagagaccacatgtatctggtggaatttgtgaccgctgctgg |
| top_var3 | aataatcatatgagagaccacatggtgctggtggaaattgtgaccgctgctgg |
| top_var4 | aataatcatatgagagaccacatggtgctggtggaatttgtgtatgctgctgg |
| top_var5 | aataatcatatgagagaccacatgtatctggtggaaattgtgaccgctgctgg |
| top_var6 | aataatcatatgagagaccacatggtgctggtggaagtggtgaccgctgctgg |
| top_var7 | aataatcatatgagagaccacatggtgctggtggaatttgtggtggctgctgg |
| top_var8 | aataatcatatgagagaccacatggtgctggtggaatttgtgtgggctgctgg |
| top_var9 | aataatcatatgagagaccacatgaccctggtggaatttgtgaccgctgctgg |
| top_var10 | aataatcatatgagagaccacatggtgctggtggaatttgtgtttgctgctgg |

**Table S3. The query sequences of the retrospective test set**. Hydrogen bonded sites are marked as pairs of single quotes (′). ‘N’ and ‘C’ represent the N-terminus and C-terminus, respectively.

| PDB ID | β-sheet Interaction | Query Sequence | PDB ID | β-sheet Interaction | Query Sequence |
|---|---|---|---|---|---|
| 2lnq | N-**LV′FF′AE′**-C<br>C-AF′FV′LK′-N | LVFFAE | 2y3l | N-**V′GG′VV′**-C<br>C-I′VV′GG′-N | VGGVV |
| 2omq | N-**L′YL′**-C<br>C-A′EV′-N | LYL | 3pzz | N-**AI′I**-C<br>C-II′A-N | AII |
| 2y2a | N-**K′LV′FF′**-C<br>C-A′FF′VL′-N | KLVFF | 3nve | N-**MH′F**-C<br>C-FH′M-N | MHF |
| 3md4 | N-**M′LG′**-C<br>C-L′MY′-N | MLG | 3loz | N-**SF′S**-C<br>C-SF′S-N | SFS |
| 4e0k | N-**WS′F**-C<br>C-FS′W-N | WSF | 6dj0 | N-**S′LT′**-C<br>C-S′VT′-N | SLT |
| 4e0l | N-**Y′LL′YY′**-C<br>C-Y′YL′LY′-N | YLLYY | 6dix | N-**V′FG′**-C<br>C-G′FV′-N | VFG |
| 4e0m | N-**I′VY′**-C<br>C-Y′VI′-N | IVY | 5e61 | N-**AI′L**-C<br>C-LI′A-N | AIL |
| 3fth | N-**LV′HS′**-C<br>C-HV′LF′-N | LVHS | 5wor | N-**V′KV′WG′**-C<br>C-I′SG′WV′-N | VKVWG |

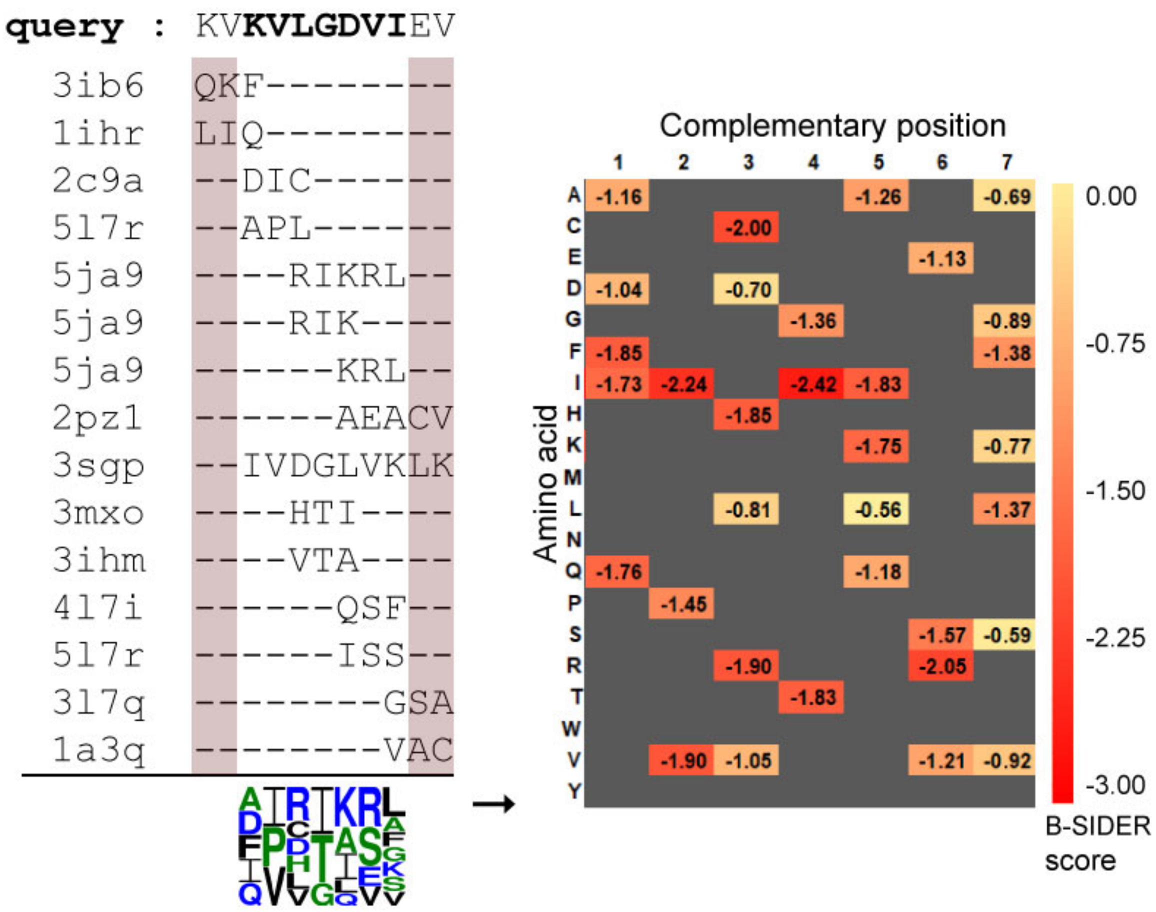


**Figure S1. Schematic description of B-SIDER scoring.** A query sequence is divided into linear fragments and they are searched against the structural database. Terminal residues can be underweighted due to the subsequence overlap. Thus, the subsequence search starts at the -2$^{th}$ position (red highlighted), if available. Complementary β-strand sequences of each match are collected and frequencies of amino acids at each position are counted.

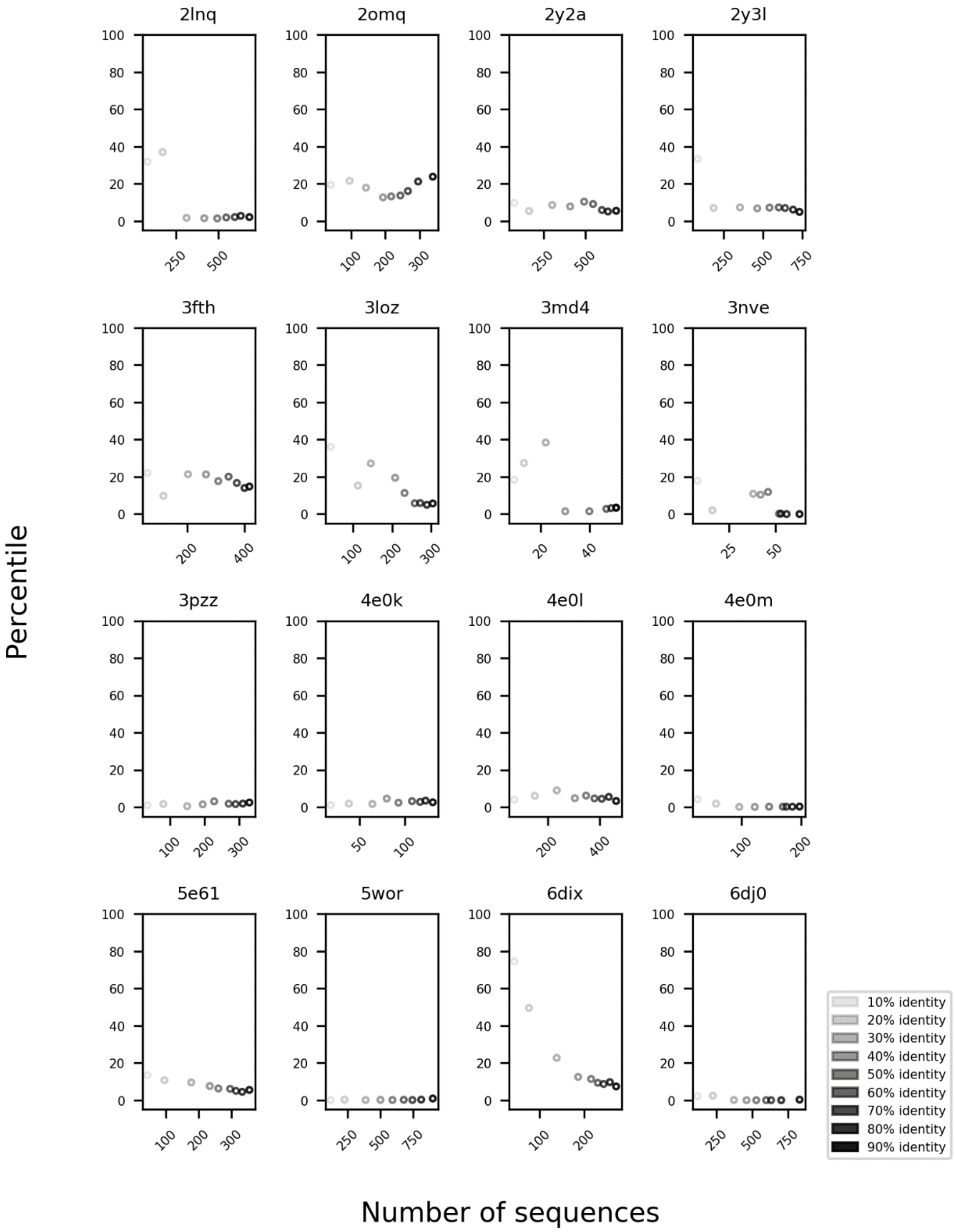


**Figure S2. Discriminatory power of the B-SIDER score at various sequence identity cutoff values (individual cases)**. The x-axis represents the number of available sequences at each sequence identity threshold. The higher the threshold is, the more sequences for generating scoring matrix.

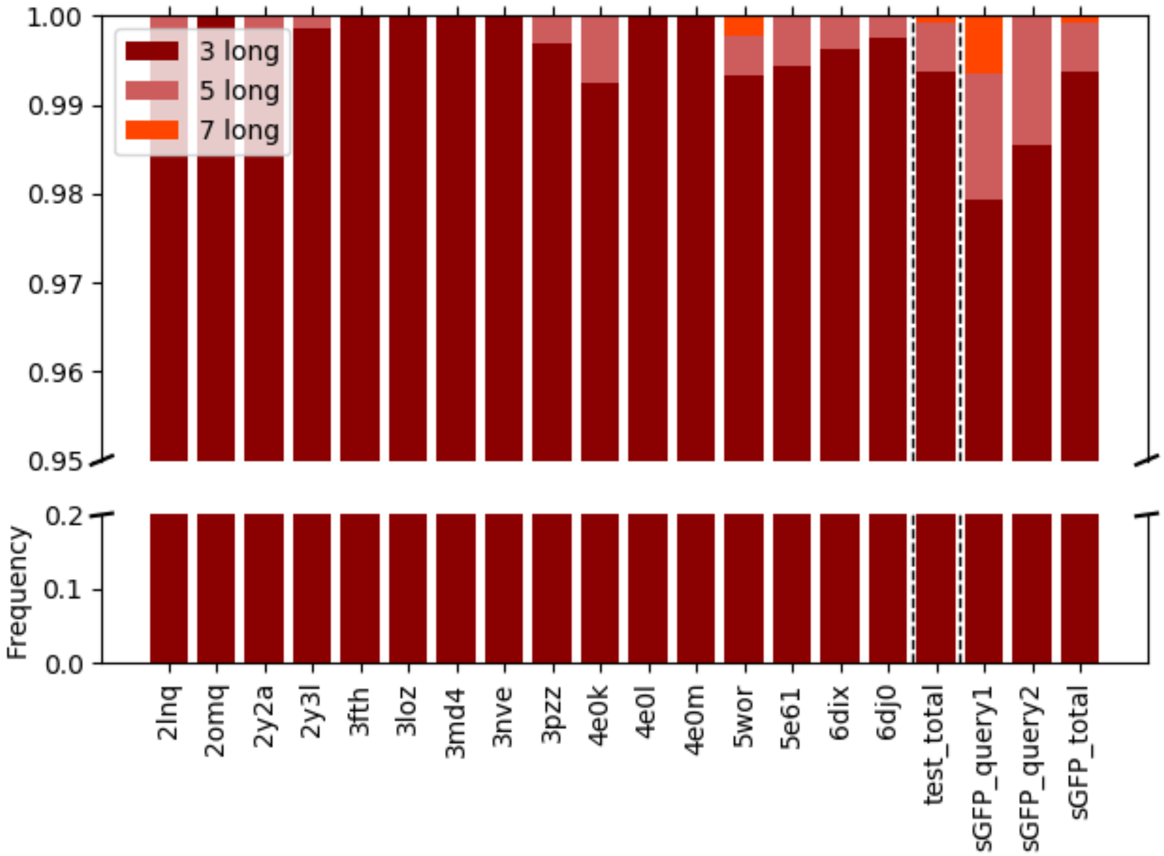


**Figure S3. Length distributions of matched subsequences.** Subsequences with three residues are predominant in all cases.

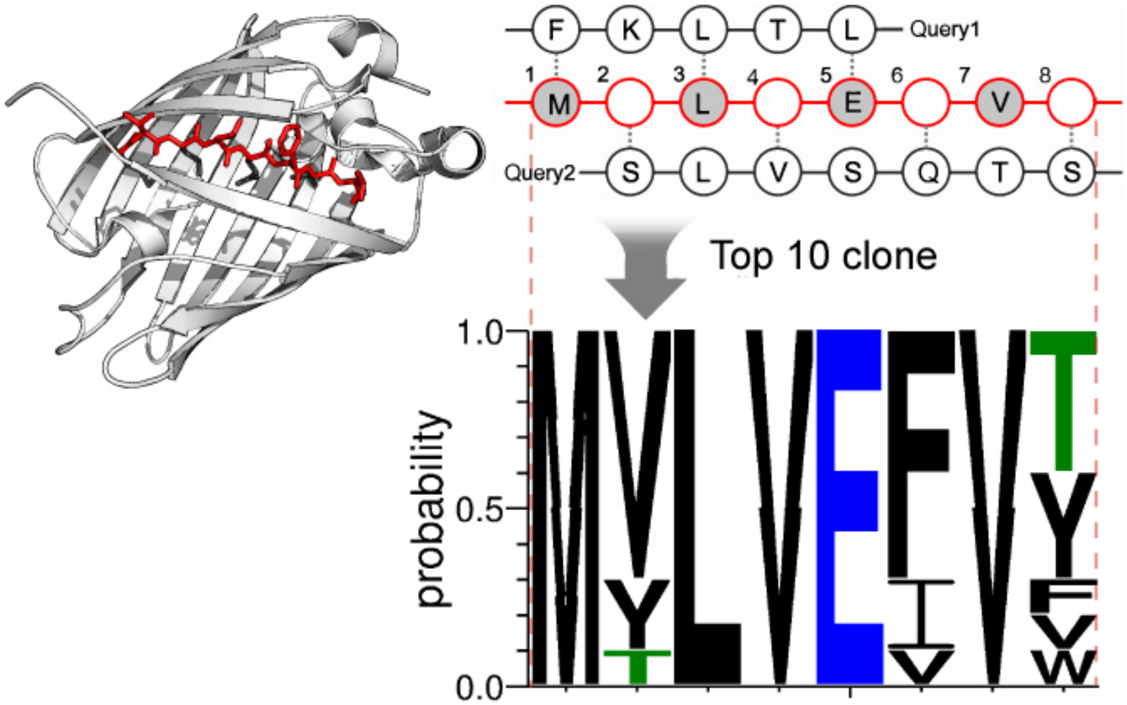


**Figure S4. The sequence logo of GFP11 variants.** The amino acid composition of the top 10 selected GFP 11 variants is presented as a sequence logo. The first three mutated positions (2,4, and 6) are highly hydrophobic, but a hydrophilic residue (threonine) is dominant at the last position.

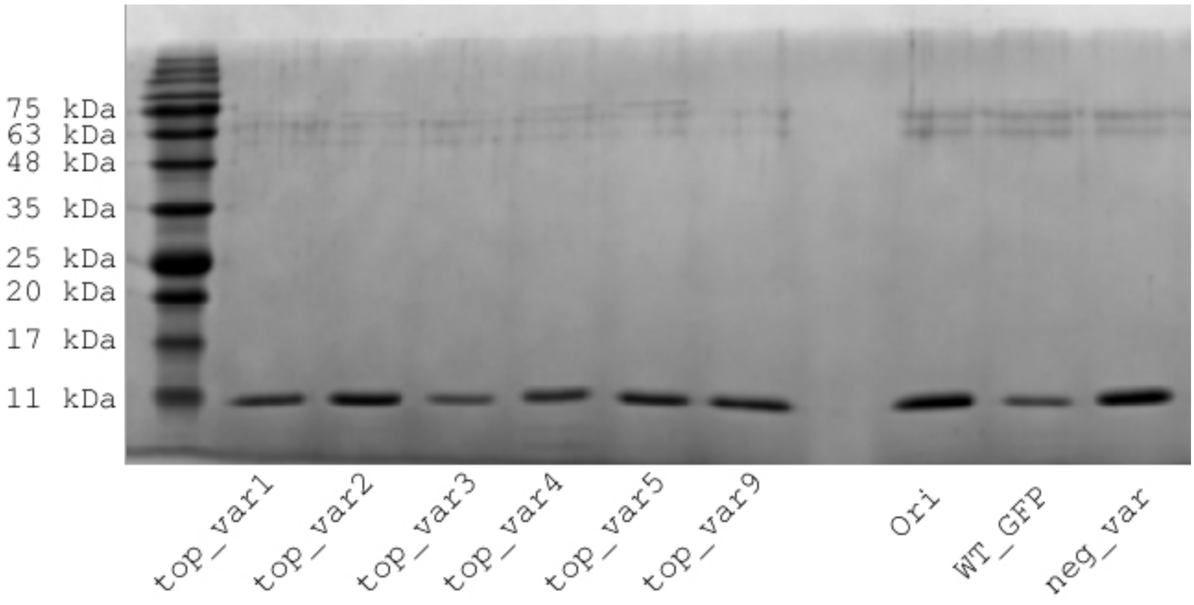


**Figure S5. SDS-PAGE analysis of GFP11 variants.** Affinity purification of designed strands was performed using Ni-NTA and desalting/buffer exchange.

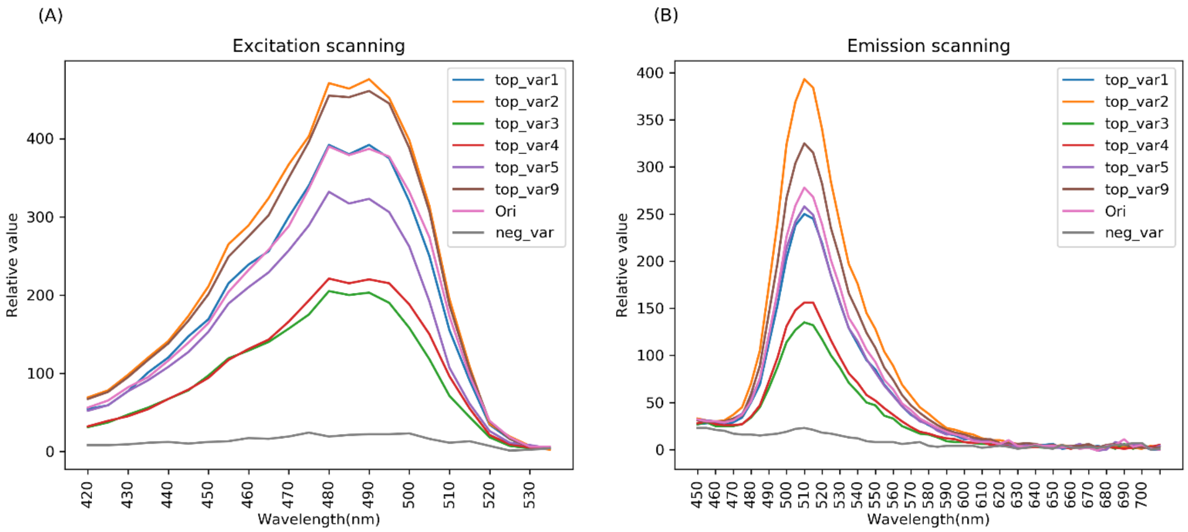


**Figure S6. Excitation/Emission spectral analyses of GFP11 variants.** (A) Scanning excitation from 420 nm to 535 nm with 5 nm interval (600 nm emission) were measured and (B) Scanning fluorescence emission from 450 nm to 710 nm with 5nm interval (400nm excitation). There are no observable differences between variants.

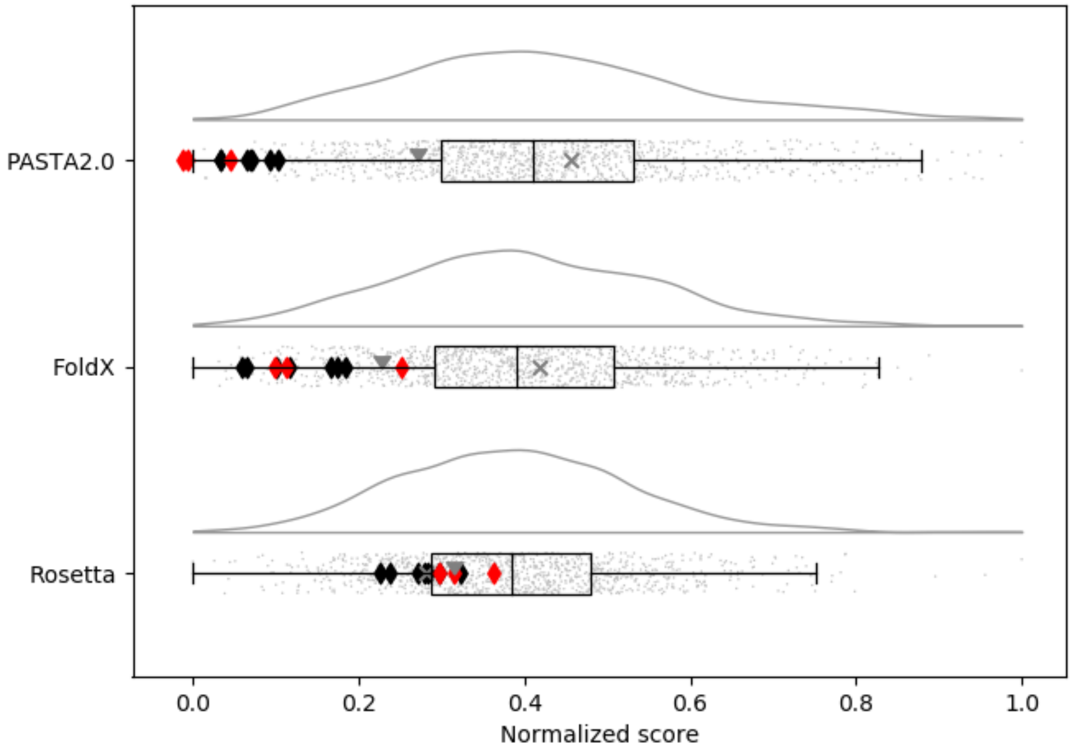


**Figure S7. Score reassessment of the GFP11 variants using PASTA2.0, FoldX and Rosetta.** The GFP11 variants generated by B-SIDER were re-assigned using other score metrics against 1,000 random sequences. All the scores were normalized for comparison. The designed GFP11 variants are designated with the diamond markers (red: stronger and faster assembled variants than the original peptide, black: the other variants than the red). The grey triangular marker and "X" represents the original peptide and the negative control, respectively, as in **Fig. 4B**.
